# Mammalian D-Cysteine controls insulin secretion in the pancreas

**DOI:** 10.1101/2022.05.17.492243

**Authors:** Robin Roychaudhuri, Timothy West, Soumyaroop Bhattacharya, Harry G. Saavedra, Lauren Albacarys, Moataz M. Gadalla, Mario Amzel, Peixin Yang, Solomon H. Snyder

## Abstract

D-amino acids are being recognized in mammals as important molecules with function. This is a first identification of endogenous D-cysteine in mammalian pancreas. D-cysteine is synthesized by serine racemase (SR) and SR^−/−^ mice produce 6-10 fold higher levels of insulin in the pancreas and plasma including higher glycogen and ketone bodies in the liver. The excess insulin is stored as amyloid in secretory vesicles and exosomes. In glucose stimulated insulin secretion studies in mouse and human islets, equimolar amount of D-cysteine showed higher inhibition of insulin secretion compared to D-serine, another closely related stereoisomer synthesized by SR. In mouse models of diabetes (STZ and NOD) and human pancreas, the diabetic state showed increased expression of D-cysteine compared to D-serine followed by increased expression of SR. SR^−/−^ mice show decreased cAMP in the pancreas followed by reduced phosphorylation of CREB (S133), lower DNA methyltransferase enzymatic and promoter activities resulting in decreased methylation of the *Ins1* promoter. D-cysteine is efficiently metabolized by D-amino acid oxidase and transported by ASCT2 and Asc1. Dietary supplementation with methyl donors restored the high insulin levels and low DNMT enzymatic activity in SR^−/−^ mice. Our data show that endogenous D-cysteine in the mammalian pancreas is a regulator of insulin secretion.

**Highlights:**

1. Serine Racemase also functions as a cysteine racemase.
2. Lack of Serine Racemase results in significantly high levels of insulin in the pancreas, plasma and larger islets.
3. D-cysteine shows greater inhibition of insulin secretion compared to D-serine.
4. Endogenous D-cysteine signals via cyclic AMP that mediates downstream CREB-DNMT1 interaction.
5. CREB-DNMT1 interaction results in hypomethylation of *Ins1* promoter that can be rescued by high methyl donor dietary supplementation rescuing high insulin levels.

## 1. Introduction

D-amino acids are mirror images of L-amino acids and their function in mammals are gradually being elucidated [2]. While much of the work in this area has focused on D-serine and D-aspartate (ligands at the NMDA receptor), little is known about endogenous mammalian D-cysteine [10]. Using novel biochemical methods (in vivo and in vitro), *Ins1* promoter bisulfite sequencing and high throughput mass spectrometric approaches, we identify endogenous D-cysteine in the mammalian pancreas and highlight its role in insulin secretion.

Our data show that mammalian D-cysteine is present in substantial amounts in the pancreas and is synthesized by enzyme serine racemase (SR). SR^−/−^ mice show reduced levels of D-cysteine and constitutively produce 6-10 fold higher levels of insulin in the pancreas and plasma and display 3 fold higher mean islet diameters. The excess insulin is stored in secretory vesicles and plasma exosomes as amyloid aggregates. Endogenous D-cysteine controls nucleotide metabolism and signals via cAMP. SR^−/−^ mice display lower levels of CREB, p-CREB (S133) and total DNMT activity in the pancreas which results in lower levels of global and *Ins1* promoter methylation. In both STZ and NOD mouse models of diabetes and human type 2 diabetes, SR expression was increased in the diabetic state with D-cysteine levels being higher than D-serine. Glucose stimulated insulin secretion in isolated mouse and human islets show D-cysteine to be more inhibitory to compared to D-serine. Endogenous D-cysteine including its metabolites and other D-amino acids may play novel roles in type 1 diabetes and in mammalian pancreatic biology.

## 2. Results

### 2.1.1 Identification of mammalian D-cysteine

We employed stereospecific bioluminescent and chromatographic methods to estimate endogenous D-cysteine in the pancreas (**Fig. 1A**). Using a bioluminescent luciferase assay (11–13) (**Fig. 1B**), we estimated levels of D-cysteine in the pancreas of mice (8). The luciferase assay is specific for D-cysteine as it involves the conjugation of cyano hydroxy benzothiazole (CHBT) with D-cysteine in presence of TCEP and base to form D-luciferin, which serves as an exclusive substrate for firefly luciferase (**Fig. 1B** adapted from Niwa et al)(1,11–14). WT pancreas contained 0.3 μmoles/g tissue while SR^−/−^ pancreas contained 0.1 μmoles/g tissue D-cysteine (**Fig. 1C**). We estimated D-cysteine levels in βTC-6 cells (β-cell line) with or without lentiviral knockdown of SR (SR#8) (**Fig. S3A**) and mouse plasma. Knock down of SR in βTC-6 cells (SR#8) elicited 2 fold reduction in D-cysteine levels (**Fig. S3B),** suggesting that SR is a biosynthetic enzyme for D-cysteine.

**Figure. 1.**
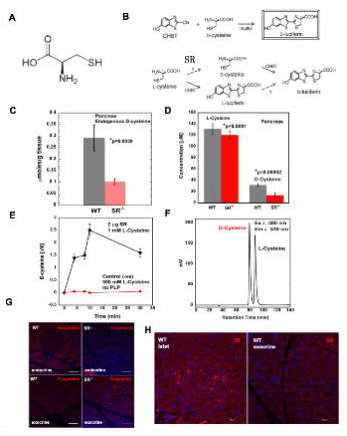

We also estimated D-cysteine levels in the pancreas using chiral HPLC following thiol labeling with ABD-F (4-(aminosulfonyl)-7-fluoro-2, 1, 3-benzoxadiazole) and fluorescent detection (**Fig. 1F**)(15,16). HPLC estimation showed four-fold molar excess of L-cysteine in the pancreas relative to D-cysteine with D-cysteine levels at approximately 30 μM in WT and 10 μM in SR^−/−^ mouse pancreas (**Fig. 1D**).

To further evaluate SR as a source for D-cysteine, we used the luciferase assay to determine racemization of 1 mM L-cysteine to D-cysteine by purified recombinant mouse SR in the presence of Mg^2+^ and PLP. Purified mouse SR efficiently racemizes L-cysteine to D-cysteine (**Fig. 1E**). Interestingly, high concentrations of L-cysteine competitively inhibit SR [19]. We used 500 mM L-cysteine in the absence of PLP as a negative control. Negative control displayed no racemization activity (**Fig. 1E**).

We next determined the localization of D-cysteine in mouse pancreas using a monoclonal antibody towards conjugated D-cysteine. D-cysteine was predominantly observed in the islets with minimal staining in exocrine pancreas (**Fig. 1G**) (**Fig. S7A**). Lack of staining in SR^−/−^ pancreas confirmed that SR is a source of D-cysteine (**Fig. 1G**). Colocalization of insulin with D-cysteine, showed D-cysteine localized mainly in β-cells (**Fig. S7B**). To correlate D-cysteine levels in the pancreas with SR, we determined expression of SR in both endocrine and exocrine pancreas. We found higher expression of SR in the islets compared to exocrine pancreas (**Fig. 1H**) [18,56].

### 2.1.2 SR controls insulin secretion and islet size

SR^−/−^ mice produced significantly higher levels of insulin in both pancreas and plasma (fed state) (**Figs. 2A, 2B**). Plasma fasting insulin levels (t=16h) were elevated in age matched SR^−/−^ mice (**Fig. S9A**). These mice also contain high levels of glycogen in the liver (**Fig. 2C**) [19]. In addition, SR^−/−^ mice produced higher levels of the ketone bodies, 2-hydroxybuyrate and 3-hydroxybutyrate in the liver (**Table. 1**) suggesting that ketone bodies may serve as a circulating energy source [20,21]. SR^−/−^ mice displayed 3 fold larger mean islet diameters and 4 fold higher total β-cell area compared to age matched controls (**Fig. S2A-B**). In tests of glucose tolerance (GTT), SR^−/−^ mice displayed a trend towards increased glucose breakdown with time (**Fig. S1A**). During the GTT, SR^−/−^ mice also displayed significantly higher levels of insulin compared to WT (**Fig. S1B**). To determine if the high insulin levels reflect abnormal transport and or processing, we measured gene expression of pre and mature insulin. Our results showed more than 10 fold higher mRNA levels of both pre and mature insulin in the pancreas of SR^−/−^ mice (**Fig. S1C**). To determine if glucose transport, islet development and insulin processing were affected, we determined their gene expression, revealing a trend towards significant upregulation in SR^−/−^ mice (**Fig. S1E-G**). Nkx6.1, a major transcription factor in pancreatic development was also upregulated (**Fig. S1D**) [22]. To assess the individual contribution of D-cysteine versus D-serine on insulin secretion, we used normal human islets, C57BL/6 mice islets and INS-1 823/13 cells which secrete insulin upon glucose stimulation.

**Figure. 2.**
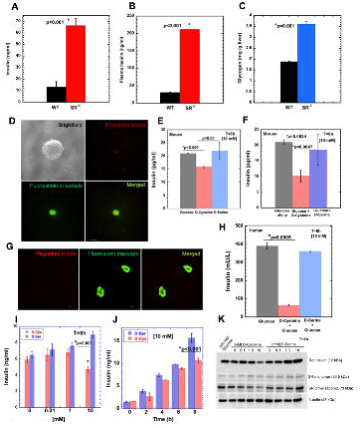

### 2.1.3 Contribution of D-cysteine and D-serine towards insulin secretion

To determine the contribution of D-cysteine and D-serine towards insulin secretion, viable islets (>95 %) from pancreas of adult C57BL/6J mice treated in culture with 10 mM concentrations of glucose, D-serine and D-cysteine individually for 6h showed significant reduction in insulin secretion in presence of D-cysteine compared to D-serine (**Fig. 2D-E**). Glucose Stimulated Insulin secretion (GSIS) performed in islets treated with 10 mM D-cysteine in presence of equimolar concentration of glucose for 6h showed significant reduction in insulin secretion compared to islets treated with equimolar concentrations of D-serine with glucose and glucose alone (**Fig. 2F**).

Normal viable (>95 %) human islets in a GSIS assay treated with 10 mM glucose, D-cysteine + glucose and D-serine + glucose for 8h at 37°C showed significantly lower levels of insulin secretion in islets treated with D-cysteine compared to D-serine (**Fig. 2G-H**, **Table 4**).

To corroborate our findings in another GSIS system, we used INS-1 cells treated with different concentrations of D-serine and D-cysteine. Our data showed similar insulin levels except at 10 mM concentration where D-cysteine was more inhibitory effect compared to D-serine (**Fig. 2I**). We next performed a time course with 10 mM concentration of D-cysteine and D-serine alone in INS-1 cells. Both stereoisomers showed similar levels of insulin secretion, however at 8 h, cells treated with 10 mM D-cysteine showed significantly lower insulin secretion compared to D-serine (**Fig. 2J**). Western blot of INS1 cells showed decrease in rat insulin at 10 mM D-cysteine compared to D-serine. This was correlated with a decrease in SR dimer for cysteine and increase for serine (**Fig. 2K**).

### 2.1.4 Excess insulin is stored as functional amyloid in secretory vesicles and exosomes

SR^−/−^ mice are fertile, healthy, have normal lifespan and produce healthy offspring. We investigated the tolerance of SR^−/−^ mice to constitutively elevated levels of insulin. A prior study showed that secretory hormones like insulin, glucagon and parathyroid hormones are stored in specialized secretory vesicles as functional amyloids [23].

We isolated secretory vesicles from pancreas of mice by differential gradient ultracentrifugation and assayed insulin containing fractions using dot blot assays [24]. Secretory vesicles from pancreas of SR^−/−^ mice contained higher amounts of amyloid compared to WT (**Fig. S4A**). To determine if these aggregates comprised insulin, we double immunostained pancreatic sections from WT and SR^−/−^ mice for amyloid and insulin [23,25,26]. Our data show increased expression and strong colocalization of amyloid and insulin in the islets of SR^−/−^ mice (**Fig. S4B**). We performed similar experiments in secretory vesicles from β-TC6 cells. Dot blot assays showed higher expression of insulin in SR#8 cells (**Fig. S4C**), in addition to increased amyloid in the secretory vesicles (**Fig. S4E**). Colocalization of amyloid and insulin in βTC-6 cells showed similar trends (**Fig. S3C**). Thioflavin-S was used to visualize amyloid aggregates in islets [27]. Results show greater Thioflavin-S staining in islets of SR^−/−^ mice (**Fig. S4I**). Since plasma of SR^−/−^ mice showed elevated levels of insulin, we isolated exosomes from plasma and assayed for size and presence of β-sheet amyloid aggregates. Electron micrographic analysis showed plasma exosomes from SR^−/−^ mice were larger compared to WT (**Figs. S4D, S4F**). ThT is a dye that intercalates individual β-sheets to produce fluorescence [28,29]. To determine if exosomes contained β-sheet aggregates, we performed a Thioflavin T (ThT) time course assay to ascertain protein assembly following lysis of exosomes and secretory vesicles. Results with plasma exosomes (**Fig. S4G**) and βTC-6 secretory vesicles (**Fig. S4H**) showed that SR^−/−^ exosomes and SR#8 vesicles produced significantly enhanced ThT fluorescence indicative of higher amounts of β-sheet aggregates. These data indicate that the high levels of insulin in SR^−/−^ mice and in SR#8 βTC-6 cells are stored as amyloid aggregates in exosomes and secretory vesicles.

### 2.1.5 D-Cysteine levels in STZ (Streptozotocin) and NOD (Non Obese Diabetes) models of Diabetes

Based on the correlation of insulin and levels of D-cysteine in the pancreas of SR^−/−^ mice and the effect of D-cysteine on mouse and human islet GSIS, we tested the levels of D-cysteine and D-serine and expression of SR in two mice models of diabetes, STZ and NOD. In an STZ model of type 1 diabetes, we injected 6-8 weeks old male C57BL/6J mice with 40 mg/kg i.p. STZ injection once daily and pancreas harvested in a time course from days 2-8 (**Fig. 3A**). Using conjugated D-cysteine and D-serine antibody, we observed increased expression of both D-serine and D-cysteine with increased duration of STZ administration (**Fig. 3B**). The expression of D-cysteine in the islets was higher compared to D-serine (**Fig. 3B**). With increased duration of STZ administration there was progressive decrease in plasma insulin levels in mice suggesting the destruction of β-cells (**Fig. S5A**) in the diabetic state that correlated with increased expression of SR (enzyme synthesizing D-serine and D-cysteine) in pancreatic lysates (**Fig. S5B**). Luciferase assay measurements of quantitative D-cysteine levels in the STZ model showed 2 fold increase in D-cysteine from 2.5 μmoles/g tissue to 4.5 μmoles/g tissue at day 8 of STZ administration (**Fig. 3F**). We also determined the levels of D-cysteine in a NOD model of diabetes (both type 1 and 2) and observed 4 fold increase in D-cysteine in diabetic NOD mice (blood glucose >250 mg/dl) compared to control NOD mice (blood glucose <120 mg/dl) (**Fig. 3G**). Expression of SR was also increased in pancreas of NOD mice correlating increased D-cysteine in diabetic NOD mice relative to control (**Fig. 3E**). To determine if the expression levels of SR, D-cysteine and D-serine in mice correlate with humans, we evaluated expression of SR in normal and type 2 diabetic human pancreas. Our data show increased expression of SR in islets of diabetic pancreas compared to normal (**Fig. 3C**). Higher expression of SR in diabetic pancreas was correlated with decreased insulin expression (**Fig. 3C**). We determined expression of D-cysteine and D-serine in normal and diabetic human pancreas and observed increased expression of both D-cysteine and D-serine in diabetic pancreas compared to normal, however the levels of D-cysteine clearly appear to be higher than D-serine in the diabetic pancreas supporting our GSIS data in mouse and human islets (**Figs. 3D and 2E, F and H**). Our data collectively show that D-cysteine has a more robust inhibitory effect on insulin secretion compared to D-serine with higher D-cysteine correlated with lower insulin levels and vice versa.

**Figure. 3.**
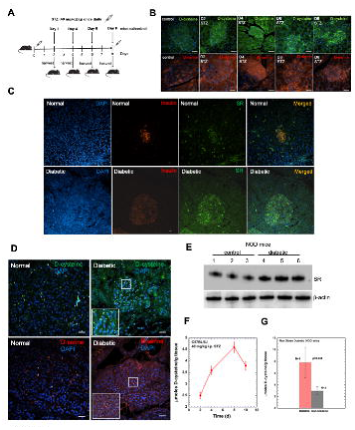

### 2.1.6 Gene Ontology and Pathway analysis

To identify proteins interacting with SR, we performed high throughput label free quantitative mass spectrometry (MS) in pancreatic lysates of WT and SR^−/−^ mice [33]. Our MS screen showed upregulation of 196 proteins (>3 fold relative to WT) (**Fig. S6A**) and downregulation of 33 proteins. Interestingly, we observed a 70 % decrease in cytosolic 10-formyl tetrahydrofolate dehydrogenase (Aldh1L1) which is involved in mitochondrial 1C metabolism and controls SAM levels (**Fig. S6B**) [34]. Protein sets obtained following enrichment analysis in ShinyGO and the Reactome database identified 560 GO categories and 134 pathways associated with translation, nucleotide binding, RNA quality control and protein transport (**Fig. 4A-B**). The 33 proteins downregulated in SR^−/−^ identified 76 GO categories and 274 pathways related to cytoskeletal proteins (intermediate filament), protein quality control, ubiquitination, proteasome and DNA damage regulation (**Fig. 4C-D**).

**Figure. 4.**
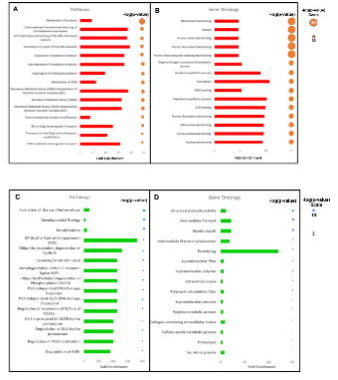

### 2.1.7 D-Cysteine signals via cAMP

We performed quantitative cAMP ELISA to determine the effect on nucleotide metabolism. Our data show significantly lower levels of cAMP (*p<0.0001 relative to WT) in the pancreas of SR^−/−^ mice compared to control (**Fig. 5A**). We also performed live cell imaging in βTC-6 cells transfected with the cAMP sensor Pink Flamindo (Pink Fluorescent cAMP indicator) and treated with equimolar amounts of L-cysteine, D-cysteine and forskolin (50 μM each) [35]. Snapshot of live cell imaging (**Movies S1-S4**) show that D-cysteine treatment produced maximal fluorescence (**Figs. 5C, 5D**). Identical experiments in HEK293 cells showed D-cysteine treatment with highest cAMP fluorescence comparable to forskolin (positive control) (**Fig. S8**). Immunohistochemistry on pancreas of WT and SR^−/−^ mice showed reduced expression of cAMP in both exocrine and islets of SR^−/−^ mice (**Fig. 5B**). Our data show that D-cysteine signals via cAMP (**Fig. 5E**).

**Figure. 5.**
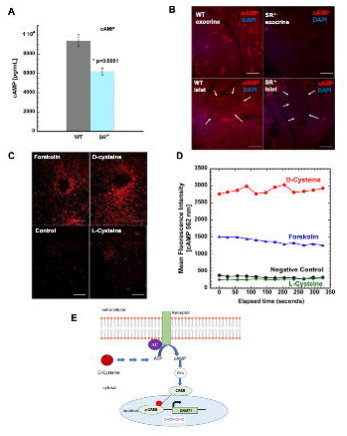

### 2.1.8 SR controls global and *Ins1* promoter methylation

To elucidate a mechanistic basis for the high levels of insulin in SR^−/−^ mice, we examined an epigenetic mode of regulation. Our rationale was based on the five-fold reduction of cytosolic enzyme 10-formyl tetrahydrofolate dehydrogenase (controls S-Adenosyl Methionine (SAM) levels and DNA methylation) in the proteomics screen (**Fig. S6B**).

To evaluate methylation of the *Ins1* promoter, we performed bisulfite sequencing around the 2500 bp region containing CpG islands (**Fig. S1H**) [36-38]. Our data showed lower levels of methylation at positions 33, 60, 113, 133, 147 and 156 of the *Ins1* promoter of SR^−/−^ pancreas with the highest difference (compared to WT) seen at positions 33, 113 and 133 (indicated by *p* values) (**Fig. 6C**). These data suggest that lower levels of methylation of the *Ins1* promoter may lead to higher insulin expression in SR^−/−^ mice. DNA dot blot assays also showed lower levels of methyl cytosine (mC) (**Fig. 6D**) and hydroxymethyl cytosine (hmC) (**Fig. 6E**) in SR^−/−^ mice.

**Figure. 6.**
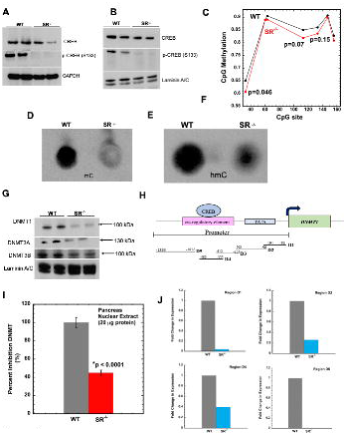

CREB is a transcription factor that controls *DNMT1* gene expression [39]. To elaborate a mechanism, we determined expression of CREB and phospho CREB. We observed lower levels of total CREB and phospho-CREB (S133) in pancreatic lysates (**Fig. 6A**) and in nuclear extracts (**Fig. 6B**).

We determined expression of DNMT1, DNMT3A and DNMT3B which were significantly reduced in SR^−/−^ mice (**Fig. 6G**). We assayed total DNMT activity in nuclear extracts of mice pancreas. Our results show an 80 % reduction in total DNMT activity in SR^−/−^ mice compared to WT (**Fig. 6I**).

To determine if lower levels of CREB affect *DNMT1* promoter activity, we performed chromatin immunoprecipitation followed by real time PCR (ChIP-qPCR). We divided the *DNMT1* promoter into 5 regions (D1-D5) spanning from −1149 to −183 bp (**Fig. 6H**) [40]. We determined fold change in CREB binding in regions D1-D5 in immunoprecipitated DNA against an input control. We detected significant reduction in CREB binding to the *DNMT1* promoter in regions D1, D2 and D5 with the exception of D3 in SR^−/−^ DNA (**Fig. 6J**). These results suggest that lower levels of total CREB and pCREB (S133) in SR^−/−^ pancreas results in lower expression of *DNMT1* and consequently decreased methylation of the *Ins1* promoter.

### 2.1.9 Methyl donor dietary supplementation rescues excess insulin secretion and DNMT activity

SAM is synthesized via the methionine cycle from multiple precursors like folate, choline, betaine and methionine (**Fig. 7A**) [34,41]. To determine if high insulin in SR^−/−^ mice could be restored to physiological levels by providing methionine, we fed WT and SR^−/−^ mice a high carbohydrate diet containing 0.8 % methionine and or a normal diet (0.4 % methionine) for 6 months (*ad libitum*). SR^−/−^ mice fed a normal diet produced 3 fold higher insulin compared to control mice. However, SR^−/−^ mice fed a high carb diet (0.8 % methionine) had significantly reduced levels of insulin compared to SR^−/−^ mice fed a normal diet (**Fig. 7B**). SR^−/−^ mice fed a normal diet had higher body weights compared to age matched WT mice (**Fig. S9B**). Plasma insulin were 8 fold lower in SR^−/−^ mice fed a high methionine diet compared to the same mice fed normal diet (**Fig. 7D**). DNA dot blot showed partial rescue with higher levels of mC and hmC in mice fed high methionine diet (**Fig. 7C**).

**Figure. 7.**
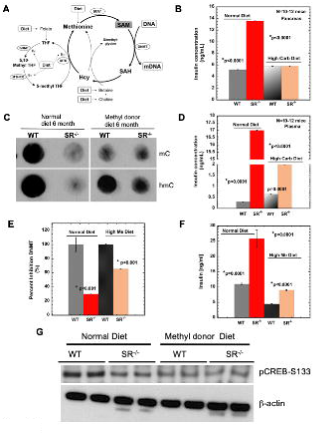

To determine if the rescue in insulin levels was due to methionine in the high carbohydrate diet, we fed age matched WT and SR^−/−^ mice an exclusive methyl donor diet containing 1.16 % methionine or a normal diet containing 0.4 % methionine for 3 months (*ad libitum*) [42,43]. Our data showed that SR^−/−^ mice fed a high methyl donor diet displayed 2.5 fold lower insulin levels compared to SR^−/−^ mice fed a normal diet (**Fig. 7F**). We next assayed total DNMT activity in pancreatic nuclear extracts from mice fed high methyl donor and or normal diet to determine rescue of DNMT activity. SR^−/−^ mice fed high methyl donor diet showed approximately two fold recovery of total DNMT activity compared to SR^−/−^ mice fed a normal diet (**Fig. 7E**). However, we did not observe rescue of CREB phosphorylation (S133) in mice fed high methyl donor diet (**Fig. 7G**). These data point to an epigenetic mode of regulation of insulin by SR that involves *Ins1* promoter methylation.

#### 2.2 D-cysteine is efficiently degraded by D-amino acid oxidase

D-amino acid oxidase (DAO) is a flavin adenine dinucleotide (FAD) dependent enzyme that regulates the physiological concentrations of various D-stereoisomers in vivo [30,31]. To determine if D-cysteine serves as a substrate for DAO, we performed in vitro kinetics with purified porcine DAO towards D-serine and D-cysteine, measuring the production of L-glutamate from ammonia and α-ketoglutarate by glutamate dehydrogenase in presence of NADPH [32]. Our kinetic data shows that porcine DAO can degrade D-cysteine equally efficiently as D-serine. DAO showed a K_m_ of 225 μM for both D-cysteine and D-serine and V_max_ of 0.002 μmoles/min and 0.0019 μmoles/min for D-cysteine and D-serine respectively. Our data indicate that D-cysteine and D-serine are both DAO substrates with similar affinities (**Table. 2**).

### 2.2.1 Endogenous D-cysteine is transported by ASCT2 and Asc1

To assess cellular transport of endogenous D-cysteine, we transfected HEK293 cells with ASCT2, a sodium dependent transporter of D-serine [44]. ASCT2 transfected HEK293 cells had significantly higher levels of intracellular D-cysteine than un transfected control (**Fig. S10A, S10E**). Similar assays with a sodium independent transporter Asc1 in HEK293 cells showed significantly higher accumulation of intracellular D-cysteine relative to control (un transfected) (**Fig. S10B, S10E**). *Ex viv*o experiments in age matched WT and Asc1^−/−^ mice (day 10 old as Asc1^−/−^ mice die by day 14) pancreas showed higher accumulation of intracellular D-cysteine in Asc1^−/−^ mice (1.8 μM) relative to Asc1^+/-^ (0.3 μM) and WT mice (**Fig. S10C**). Pancreatic lysates of WT and Asc1^−/−^ mice showed reduced SR expression in liver and pancreas (**Fig. S10D**). High levels of D-cysteine (> 1mM) inhibit racemization activity of SR [17]. Our data suggests that high levels of intracellular D-cysteine may inhibit expression and racemization by SR in the liver and pancreas.

## 3. Discussion

Our work is the first report on endogenous D-cysteine synthesized by SR in mammalian pancreas. We estimated endogenous D-cysteine using a novel, stereospecific bioluminescent luciferase assay and chiral HPLC [12]. Cysteine is the fastest in vitro racemizing amino acid with a t_1/2_ of 2 days highlighting its tight regulation in vivo [46]. Racemization serves to effectively regulate its tissue levels that may be dependent on the cellular microenvironment and or its function during development. Evidence for L-cysteine being a substrate for SR was observed in a study of L-serine racemization where cysteine showed more than 95 % inhibition of L-serine racemization [47]. Based on our kinetic data and prior work, SR also functions as a cysteine racemase [11]. Dual substrate specificity of SR implies that it regulates concentrations of both D-serine and D-cysteine. Prior work has suggested different temporal profiles for D-cysteine and D-serine during neural development implying discrete functions [10,11].

High insulin levels in SR^−/−^ mice are constitutive and correlated with significantly increased islet size, higher body weights including higher glycogen and ketone bodies in the liver. Interestingly, glucagon expression remained unchanged. Hypoglycemia due to high insulin levels have been shown to generate insulin antibodies that slow glucagon secretion and plasma glucose recovery [48]. Higher ketone bodies in SR^−/−^ mice imply utilization of alternate sources of energy under conditions of glucose depletion. To address the individual contribution of D-cysteine versus D-serine on insulin secretion, we utilized normal human and C57BL/6J mouse islets including INS-1 832/13 cells that secrete insulin in response to glucose and other secretagogues (57). D-cysteine showed a more potent effect at inhibiting insulin secretion compared to D-serine in absence of glucose in both islets and INS-1 cells. In GSIS assays, the individual effects of D-serine and D-cysteine on inhibition of time dependent insulin secretion by glucose were observed with D-cysteine having a more potent effect compared to D-serine [58].

SR is expressed in islets [56, 60], however its role in diabetes is unknown. Prior work by Ndiaye et al implicate SR as a type 2 diabetes susceptibility gene involved in beta cell function [61]. Our data in the STZ (type 1) and NOD (both type 1 and 2) models of diabetes including human type 2 diabetic pancreas show increased SR and D-cysteine in the diabetic state implicating regulation of insulin secretion. Contribution of D-cysteine versus D-serine was studied in isolated mouse and human islets including INS-1 832 cells and showed marked inhibitory effect on insulin secretion by D-cysteine compared to D-serine arguing for differential insulin regulatory effects by both stereoisomers.

To elucidate the tolerance of SR^−/−^ mice to constitutively high levels of insulin, we investigated its storage in vivo. Colocalization studies in mice and βTC-6 cells show that insulin is stored in secretory vesicles and plasma exosomes as β-sheet aggregates. Storage of hormones as functional amyloid serves a regulatory function [23].

One-carbon metabolism by the folate and methionine cycles integrates amino acids, glucose and vitamins for synthesis of lipids, nucleotides and proteins including substrate for methylation reactions. Based on this rationale, we estimated levels of cAMP and observed significant reduction in SR^−/−^ pancreas. Live cell imaging monitoring real time dynamics of cAMP showed that stimulation of both HEK293 and βTC-6 cells with D-cysteine resulted in enhanced cAMP activation.

We performed mass-spectrometry based proteomics to identify proteins under regulatory control of D-cysteine and D-serine. Our MS screen found >2 fold upregulation of islet regenerating factor, proteins involved in transport, ribosomal synthesis, ubiquitination and chaperones in SR^−/−^ pancreas. Interestingly, cytosolic 10-formyl tetrahydrofolate dehydrogenase, which is involved in mitochondrial 1C metabolism and controls SAM levels was downregulated 5 fold. Based on this observation, we investigated DNA methylation and observed decrease in mC and hmC DNA modifications in SR^−/−^ pancreas in addition to lower methylation of the *Ins1* promoter. Our data suggests that hypomethylation of the *Ins1* promoter may serve as a plausible explanation for higher insulin levels in SR^−/−^ mice. GO and pathway enrichment analysis reveal that SR and or its product may play roles in cellular metabolism, cytoskeleton assembly, folate biosynthesis and mitochondrial electron transfer, implying novel functions in the pancreas.

CREB is a transcription factor that controls *DNMT1* expression [39]. To elucidate a mechanism, we determined levels of total and phospho CREB (S133) and observed lower levels of both in SR^−/−^ pancreatic cell and nuclear extracts. Expression of DNMT1, 3A and 3B enzymes were decreased in SR^−/−^ pancreas, resulting in reduced total DNMT enzymatic activity. ChIP-qPCR results show that SR^−/−^ pancreas have lower *DNMT1* promoter activity due to lower levels of CREB occupancy at the *DNMT1* promoter resulting in lower expression and activity. High glucose concentrations in rat INS-1 cell line and Zucker diabetic fatty rats showed increase in *DNMT1* mRNA resulting in increased DNA methylation of *Ins1* promoter and suppression of *Ins1* mRNA expression [50]. cAMP also has a regulatory effect on the transcriptome by enhancing DNA hydroxymethyation via augmentation of intracellular Fe(II) pool [51].

To test our DNA methylation hypothesis, we performed two dietary rescue experiments providing higher methionine as a source of methyl donors [52]. Our 6 months high carbohydrate and high methyl donor supplementation led to a significant decrease in insulin levels both in the pancreas and plasma of SR^−/−^ mice. A 3 month experiment with high methyl donors (1.18% methionine versus 0.8 % in the high carb diet) was equally effective, resulting in 3 fold reduction of pancreatic insulin levels. We observed partial rescue of total DNMT activity and mC and hmC levels in mice fed a high methyl donor diet. Collectively, our results demonstrate that SR controls methylation of the *Ins1* promoter in the pancreas.

Transport of D-amino acids regulates its cellular concentration in vivo. Asc-1^−/−^ mice die by postnatal day 14. Endocrine and exocrine pancreatic cell fate determinants are established by E14.5 [53]. Transport experiments performed on ex vivo pancreas from viable day 10 Asc-1^−/−^ mice and in HEK293 cells show increased accumulation of intracellular D-cysteine, suggesting that Asc1 transports D-cysteine. Increased D-cysteine accumulation also resulted in reduction in expression of SR in the liver and pancreas of Asc-1^−/−^ mice, suggesting a feedback inhibition that may regulate intracellular D-cysteine. Our studies in a simplified HEK293 cell culture show Asc-1 and ASCT2 as transporters of D-cysteine.

DAO regulates the concentrations of various D-amino acids *in vivo*. Our kinetic data showed DAO with similar affinities towards D-cysteine and D-serine. A prior study showed that competition assays between D-serine and D-cysteine had no effect on the activity of DAO, supporting the idea that D-cysteine may be a marginally better substrate then D-serine [14].

A limitation of our study is the use of mice with germline deletion of SR. Our attempts at conditional deletion of SR with *Pdx-1* and *Ins1* cre reporter mice did not yield satisfactory cre expression and recombination in the developing pancreas and insulin secreting β-cells respectively. While *Pdx-1* cre deletion show leaky expression in the brain [54], *Ins-1* cre deletion is subject to suppression by extensive DNA methylation, which is the basis of high insulin in SR^−/−^ mice [55].

Cysteine due to its redox activity has additional isomers [12]. Their D stereoisomers could potently regulate insulin levels with novel roles in pancreatic development. Inhibitors of SR, and D-cysteine may have therapeutic implications for glucose homeostasis and possibly in type 1 diabetes.

## 4. Conclusions

We show that endogenous mammalian D-cysteine in the pancreas, is synthesized by SR and regulates insulin secretion. Reduced cAMP due to deletion of SR leads to decreased phosphorylation of CREB resulting in decreased activity of *DNMT1* promoter and enzymatic activity, which may be responsible for reduced methylation of the *Ins1* promoter, affecting insulin synthesis and secretion.

## 5. Materials and Methods

### Reagents and Antibodies

Purified L-cysteine, D-cysteine, L-serine, D-serine, CHBT (cyano hydroxy benzothiazole) were purchased from Sigma Chemical Corp. TCEP HCl (Tris (2-carboxy ethyl phosphine) hydrochloride) was purchased from Thermo Scientific. All salts for buffers and reagents were of research grade and high purity. Milli Q water was used to make buffers and solutions for experiments. HPLC grade solvents and water (Fisher Scientific) was used for HPLC estimation of D and L-Cysteine. Anti-conjugated D-cysteine antibody (AB-T050; Advanced Targeting Systems, San Diego, CA). Anti cAMP mouse monoclonal antibody (MAB2146-SP; Novus Biologicals). Rabbit anti DNMT1antibody (Aviva; ARP37033_P050), mouse anti DNMT3A (PCRP-DSHB Iowa), rabbit anti DNMT3B antibody (GeneTex; GTX 129127), rabbit anti CREB (CST; 48H2), rabbit anti phospho-CREB (S133) (CST; 87G3), rabbit anti SR antibody (CST; D5V9Z), rabbit anti SLC7A10 polyclonal antibody (G-Biosciences; ITA8971).

### Mice

Male C57BL/6J 8-12 weeks age was purchased from Jackson Labs (strain #000664). SR^−/−^ (Serine Racemase knock out) mice were a property of Dr. Solomon H. Snyder at Johns Hopkins University School of Medicine and maintained in house. All mice were maintained as per IACUC rules and regulations at Johns Hopkins School of Medicine and the University of Maryland School of Medicine.

### Insulin ELISA

Quantitative estimation of insulin from plasma and pancreas was performed using the Ultra Sensitive Mouse Insulin ELISA Kit (Cat# 90080; Crystal Chem). The protocol was followed exactly as mentioned for the high range assay. Standard curve for insulin was developed prior to the actual experiment (high range). Each individual sample was done in quadruplicate. For treatment groups, the samples were pooled in their respective groups and analyzed. The absorbance was read at 450 nm in a 96 well plate reader and also at 630 nm and data plotted after correction at 630 nm.

### Analysis of Gene expression

RNA was isolated from pancreas, islets and acinar cells from WT and SR^−/−^ mice using RNeasy Mini kit (Qiagen). Gene expression studies related to glucose homeostasis and pancreas development were performed using the SYBR green method. cDNA synthesis was performed using High Capacity cDNA synthesis kit (Applied Biosystems). 100 ng of synthesized cDNA was used in each well of a 96 well 0.1 ml MicroAmp PCR plate (Applied Biosystems) along with SYBR Green master mix, nuclease free water and gene specific primers (10 μM final concentration). 18S rRNA was used as a housekeeping gene and non-template control was added in each plate as a control. The PCR was run on an ABI Step One Plus instrument. The raw data obtained was analyzed on StepOne data analysis software (StepOne Plus) and mean C_T_ values obtained. ΔC_T_ and ΔΔC_T_ were calculated and fold change in gene expression obtained from 2^exp-ΔΔCT^. Primer sequences are listed in Table.3.

### Glucose Tolerance Tests

Glucose tolerance test (GTT) was performed on a 16 h fasted WT and SR^−/−^ age matched mice injected i.p with D-glucose (2 g/kg body weight). Blood glucose level was monitored by tail bleeding immediately before and at indicated times after injection using Contour glucometer (Bayer Co, Japan) and Contour blood glucose test strips (Cat#7097C; Ascensia Diabetes care Inc, NJ). Blood glucose measurements were obtained from tail veins at indicated time points post injection. A small drop of blood from the tail was placed on a new glucose strip each time, inserted into the glucometer and value recorded.

### Plasma fasting Insulin

Age matched WT and SR^−/−^ mice (8-12 weeks old) were fasted for 16 h with only access to water. Mice were euthanized and blood obtained by cardiac puncture using a 1 ml tuberculin syringe (BD Biosciences). The harvested blood was stored in a BD Microtainer tube (cat# 365985) with lithium heparin on ice. Plasma was obtained by centrifuging the microtainer tubes at 3000 rpm for 15 minutes at 4°C. After centrifugation the cell free supernatant layer (plasma) from the top was carefully removed into a tube containing protease inhibitors (1:20 v/v) and kept on ice. The isolated plasma was assayed for insulin using mouse insulin ELISA as mentioned. Plasma insulin levels during the GTT were assayed in an exactly similar fashion after administering the required dose of glucose. Plasma for each genotype was pooled from N=5-8 mice and assayed in triplicate or quadruplicate in a 96 well format. Data were subtracted from blank wells containing sample diluent alone. Error bars obtained refer to SD. The experiment was independently performed 3 times.

### Glycogen Estimation

Glycogen was estimated from age matched WT and SR^−/−^ mouse liver. A small portion of the liver (approx. 0.5-1 g wet tissue) was dissected and minced in a petri dish and dissolved in 4 times the volume (w/v) of 10 % perchloric acid and homogenized in a handheld homogenizer. Following homogenization, the sample was centrifuged at 300 g for 10 minutes. A portion of the supernate was extracted with 1:1 (v/v) ethanol and centrifuged at 1700 g for 10 minutes at 4°C. The supernate was discarded and the pellet resuspended in the same volume of water as the initial volume of supernate (approx. 200 μl). Fifteen μl of the supernate was removed for assay in a 96 well plate. The sample was diluted in 15 μl water followed by addition of 30 μl 6.5 % phenol and 150 μl of 85 % sulfuric acid in a total volume of 210 μl. The samples were run with an N=5 replicates. Blank wells contained water instead of sample. The plate was read in a 96 well plate reader at 420 nm and the amount of glycogen quantified based on a standard curve [21].

### Estimation of ketone bodies by NMR

Ketone bodies extraction and estimation from livers of WT and SR^−/−^ mice was performed using ^1^H NMR at the Metabolomics Innovation Centre (TMIC, Canada). Briefly, samples were extracted and prepared as mentioned in [22]. The tissue was homogenized and fractionated to obtain a biphasic mixture. The upper polar phase (water-soluble metabolites) and lower non-polar phase (lipid-soluble metabolites) were carefully separated. The water-soluble tissue extract was transferred to a 1.5 ml eppendorf tube, to which an additional standard NMR buffer solution was added. The buffer consisted of 750.0 mM potassium phosphate (pH 7.0), 5.0 mM 2,2-dimethyl-2-silapentane-5 sulfonate (DSS-d6), 5.84 mM 2-chloropyrimidine-5-carboxylic acid, and D_2_O (54 % v/v in H_2_O). All ^1^H-NMR spectra were collected on a Bruker Avance III Ascend 700 MHz spectrometer equipped with a 5 mm cryo-probe (Bruker Biospin, Rheinstetten, Germany). ^1^H-NMR spectra were collected at 25°C using the first transient of a noesy-pre saturation pulse sequence. This pulse sequence was selected based on its excellent reproducibility and quantitative accuracy. NMR spectra were acquired with 128 scans employing a 4 second acquisition time and a 1 second recycle delay. Prior to spectral deconvolution, all free induction decays (FIDs) were zero-filled to 240,000 data points and a 0.5 Hz line broadening function was applied. The methyl singlet of the added DSS (set to 0.00 ppm) served both as an internal chemical shift referencing standard and as an internal standard for quantification. All ^1^H-NMR spectra were processed using the Chenomx NMR Suite 8.1 software package (Chenomx Inc., Edmonton, Canada) for compound identification and quantification as described [22].

### Quantitative proteomic analysis by nano LC-MS/MS

Global proteomic analysis of pancreatic lysates from WT and SR^−/−^ mice was performed on a nano LC-MS/MS platform by Creative Proteomics (45-1 Ramsey Road, Shirley, NY 11967, USA). The tissues were lysed in lysis buffer containing 8M urea with 1% protease inhibitor and 1% phosphatase inhibitor by sonication. Samples were centrifuged at 12,000 g for 10 minutes at 4°C and supernates harvested. Protein concentration was estimated using BCA kit. Protein digestion was performed after reduction (by 10 mM DTT) and alkylation (by 20 mM iodoacetamide) by trypsin at 37°C with overnight incubation at a trypsin: sample ratio of 1:50. Samples were centrifuged at 12,000 g and 4°C for 10 minutes. The sample was lyophilized and resuspended in 20 μl of 0.1% formic acid before LC-MS/MS. LC-MS/MS was performed on Ultimate 3000 nano UHPLC system (ThermoFisher Scientific, USA) with an analytical PepMap C_18_ column. Mobile phase A was 0.1% formic acid in water and B was 0.1% formic acid in 80 % acetonitrile. Flow rate was 250 nL/min. MS was performed on an Orbitrap Q HF mass spectrometer (ThermoFisher Scientific, USA) with a spray voltage of 2.2 kV and capillary temperature of 270°C. MS resolution was 60000 at 400 m/z and MS precursor m/z range was 300.0-1650.0. For data dependent MS/MS, up to top 20 most intense peptide ions from the preview scan in Orbitrap was selected. Data analysis following LC/MS-MS was performed using Maxquant (1.6.2.6) against a mouse protein database. The protein modifications were carbamido methylation (C) (fixed), oxidation (M) (variable); enzyme specificity was set to trypsin; maximum missed cleavages were set to 2; the precursor ion mass tolerance was set to 10 ppm, and MS/MS tolerance was 0.6 Da. Only highly confident identified peptides were chosen for downstream protein identification analysis [40].

### DAO (D-amino acid oxidase) assay

Equimolar amounts of D-cysteine and D-serine (20-500 μM) was prepared in water and diluted in 100 mM Tris buffer pH 7.5 and added to 0.026 μM (0.2 μg protein in 200 μl total reaction volume) purified porcine DAO in 100 mM Tris buffer pH 7.5, 100 μM FAD, 100 μM NADPH followed by 1 μg glutamate dehydrogenase [38]. The assay for each substrate was performed separately in a 96 well black clear bottom plate. The plate was incubated at 37°C for the duration of the assay and read at t=0, 2, 4, 6, 8, 10, 15, 20, 25, 30, 40, 50 and 60 minutes at 340 nm. The decrease in absorbance was followed and data plotted to obtain the kinetic parameters.

### Purification of mouse Serine Racemase (SR)

Mouse SR (pcDNA3.1 His tag expression vector) was expressed and induced in BL21 (DE3) competent *E. coli*. A colony was inoculated into 100 ml of LB medium with 50 ug/ml of kanamycin for 18 hrs. Later, the cells were expanded to an 8 L culture. When the OD reached 0.4, 0.2 mM ITPG was added and the cells were grown at 18°C, 200 rpm for 18 h. Cells were centrifuged at 10,000g and the pellet was frozen at −80°C. Cells were lysed with a microfluidizer and lysis buffer (50 mM Tris pH 8.0 containing 150 mM NaCl + 1 mM TCEP and 15 μM PLP). The supernatant was passed through a 5 ml Ni^2+^ affinity column (GE Healthcare) and eluted with the same lysis buffer containing imidazole. The protein eluted at about 200-250 mM imidazole. The relevant fractions were identified by SDS-PAGE, pooled and concentrated to 10 ml and applied to a 320 ml size exclusion column (HiLoad ® 26/600 Superdex ® 200, GE Healthcare). The column was equilibrated with 30 mM PIPES pH 7.4 buffer containing 100 mM NaCl + 1 mM TCEP + 15 μM PLP and eluted with the same buffer. Fractions of 8 ml were collected in an automated AKTA FPLC system (GE Healthcare) and purity was determined with SDS-PAGE. Purified mouse SR was concentrated using a Pierce protein concentrator (Thermo Scientific) 10K molecular weight cut off filter. The concentration was determined using a UV spectrometer and an extinction coefficient of 31775 M^-1^ cm^-1^. The concentrate was assayed for amount of protein and used in enzyme kinetic assays.

### Serine racemase luciferase assay

Racemization of L-cysteine to D-cysteine was measured using the luciferase assay for D-cysteine (1,14–16). Purified serine racemase (2 μg protein) was incubated at 37°C with 50 μM PLP, 1 mM Mg^2+^ and 100 mM Tris-Cl pH 7.8 buffer along with 200 μM L-cysteine as substrate in a total volume of 600 μl. Concentrations of reagents mentioned are final concentration in the assay. At 5, 10, 15, 30 and 60 minutes, an aliquot of the reaction mixture (100 μl) was removed and subjected to the luciferase assay (as mentioned above) to measure the amount of D-cysteine produced using bioluminescence. The amount of D-cysteine (estimated using a standard curve) was plotted versus time.

### Immunohistochemistry

Immunohistochemical experiments were performed on WT and SR^−/−^ mice pancreas. Briefly, the mice were euthanized by CO_2_ narcosis and the pancreas removed under a dissecting microscope and incubated in 4 % paraformaldehyde in PBS at 4°C for 48 h with gentle shaking. The tissue was paraffin embedded and sectioned at the Johns Hopkins Oncology Core Services. The sections were de paraffinized using Histoclear followed by a series of 5 min washes in absolute alcohol, 95% alcohol and PBS. Permeabilization of sections was done in 0.5% Triton X-100 in PBS for 20 min. Antigen retrieval was done in 10 mM sodium citrate buffer containing 0.05 % Tween 20 pH 6.0 by boiling sections for 1 min in the microwave and then cooling at room temperature (RT) for 20 min. Primary antibody dilutions were at 1:500 in 1% BSA in PBS and secondary antibody dilutions were at 1:1000 in the same buffer. The sections were washed twice in PBS after incubation with primary and secondary antibodies. DAPI contained in the mounting medium was used for nuclear staining. The slides were imaged using spinning disk confocal microscopy.

### Global DNA mC and hmC analysis

To determine global methylation (mC) and hydroxymethylation (hmC) of cytosine in DNA, dot blot was performed on DNA from pancreas of age matched WT and SR^−/−^ mice. DNA was isolated using Pure Link Genomic DNA isolation kit (Invitrogen; ThermoFisher Scientific) and quantified using a nanodrop. Nitrocellulose (NC) membrane was wet in 20X SSC buffer at RT for 10 minutes along with a same size Whatman filter paper. The prewet NC membrane along with the filter paper was placed in a spot blot (Schleicher & Schuell, Keene, NH) micro sample filtration manifold. Five hundred μl of 20X SSC buffer was added to each well in which the sample would be added and allowed to drain under suction. DNA samples (200 ng) were prepared for application to the slot by adding 20X SSC buffer to give a final concentration of 6X SSC in a total volume of 200 μl. The DNA was denatured by heating it at 100°C for 10 min and placed on ice. An equal volume of 20X SSC buffer was added to the sample and mixed. The DNA sample was centrifuged for 5 sec and then applied to the prewet wells and allowed to drain completely till the membrane was dry. The NC membrane was removed from the manifold and incubated in a petri dish containing filter paper soaked in denaturation solution (1.5M NaCl + 0.5 M NaOH) for 10 min. The membrane was next transferred to a similar petri dish containing neutralization solution (1.0 M NaCl + 0.5 M Tris-Cl pH 7.0) and incubated for 5 min. After incubation, the membrane was placed on a filter paper prewet with 20X SSC buffer with DNA side up and crosslinked using a Stratalinker UV Crosslinker in the auto crosslink mode (preset exposure of 1200 microjoules x100). The crosslinking time was 25-50 seconds. The crosslinked DNA on the NC membrane was placed in blocking buffer (5% BSA (w/v) in TBS-T) and incubated at RT for 30 min on a shaker. Primary antibodies to mC (Cat# A-1014; Epigentek Inc) and hmC (Cat# A-1018; Epigentek Inc) were added separately to the membranes at 1:2000 dilution in blocking buffer and incubated at 4°C overnight with gentle shaking. The membranes were washed 4 times 15 min each at RT with shaking in TBS-T following which they were incubated with respective secondary antibodies at 1:5000 dilution in blocking buffer for 60 min at RT. The membranes were washed 4 times as mentioned above and developed using ECL reagent (Super Signal West Pico Plus) for 5 min and membranes developed on an X-ray film.

### Targeted Bisulfite sequencing of Insulin promoter

Gene-specific DNA methylation was performed by CD Genomics Co. (Shirley, NY) on DNA samples from pancreas of age matched WT and SR^−/−^ mice. DNA methylation was assessed by a next generation sequencing based bisulfite sequencing PCR (BSP) according to previously published methods (43–45). BSP primers were designed using the online MethPrimer software. Genomic DNA (1 μg) was converted using the ZYMO EZ DNA Methylation-Gold kit (ZYMO) and one twentieth of the elution products were used as templates for PCR amplification. For each sample, BSP products of multiple genes were generated, pooled equally and subjected to adapter ligation. Barcoded libraries from all samples were sequenced on the Illumina Hiseq platform using paired-end 1150 bp strategy. After the preparation of the library, Qubit 2.0 and Agilent 2100 were used respectively to detect concentration of the library and insert size. Effective concentration (> 2 nM) of the library was quantitatively determined by qPCR to ensure library quality. The original data obtained from the high throughput sequencing platforms were transformed to sequenced reads by base calling. The level of methylation at each site of the insulin promoter between sample groups were statistically analyzed based on the methylation level of each site and data plotted with *p* values for N=2 samples per genotype.

### Dietary supplementation studies

8 weeks old age matched WT and SR^−/−^ mice were fed 70% carbohydrate diet containing 0.8 % methionine (Teklad Custom Diet TD.98090, Envigo, WI) for 6 months (*ad libitum*). Sufficient quantity of food was maintained in the cages and replaced every week for the entire duration of the experiment. Control mice were fed normal chow (Teklad Global 18% Protein Rodent Diet, 2018S, Envigo, WI) containing 0.6 % methionine. The normal diet was replaced at regular intervals by the animal care staff as per IACUC regulations. After completion of 6 months, blood glucose was measured and subsequently the animals euthanized by CO_2_ narcosis. Plasma was isolated by cardiac puncture and harvested using a butterfly needle attached to a 1ml syringe. The blood was collected in a lithium heparinized tube (BD Microtainer; REF 365965) and placed on ice. The tubes were centrifuged at 3000 rpm for 15 minutes at 4°C and the supernatant isolated in a separate tube containing proteinase inhibitors (Sigma Protease Inhibitor Cocktail). In addition to plasma, the pancreas was removed for ELISA and mC (methyl cytosine) and hmC (hydroxymethyl cytosine) based DNA dot blot analysis. All groups had a minimum of 12-15 mice per group.

In a separate experiment (3 months), 8 weeks old age matched WT and SR^−/−^ mice were fed a 3X methyl supplemented (3MS) diet containing 17.3 g choline, 15 g betaine, 7.5 g methionine (1.15 % methionine), 0.015 g folic acid, 1.5 % Vit B_12_ and 0.661 g zinc sulfate heptahydrate (Teklad Custom Diet T.110835, Envigo, WI) for 3 months along with age matched control mice that were fed normal chow for the same length of time (*ad libitum*) [49]. After completion of 3 months, similar parameters as above were measured to determine rescue. All groups had a minimum of 12-15 mice.

### Separation of L and D cysteine

Purified L and D cysteine were separated using the Astec Chirobiotic T chiral HPLC column (5 μm particle size, L x 1.D. 25 cm x 4.6 mm) (Suppelco) attached to a Waters 2690 alliance separations module connected to a fluorescence detector with 80-20 % (v/v) 20 mM ammonium acetate-methanol solvent by isocratic elution. The flow rate of the solvent was 0.1 ml/min. The eluates were detected by fluorescence with excitation λ 380 nm and emission λ 510 nm. The column was maintained at room temperature. Both L and D Cysteine from liver and pancreatic homogenates were extracted by homogenization in 100 mM HEPES buffer pH 7.5 using a handheld tissue homogenizer on ice with 10-15 strokes. The homogenates were then lysed using a sonicator three times with a pulse of 10-15 sec per pulse on ice. The samples were then centrifuged at 16,000 g for 30 min at 4°C. After centrifugation the supernate was harvested in a pre-chilled tube. Proteins in the supernates were precipitated by adding 12 N HCl to the lysate to give a final concentration of 2 N HCl. The sample was cooled on ice for 30 min and centrifuged at 16,000 g for 30 min at 4°C. The supernate was harvested that contained the stereoisomers of cysteine. The supernatant was neutralized by adding one equivalent NaOH (v/v) of 10 N NaOH followed by addition of 1 volume equivalent (v/v) of 1 mM ABDF (4-(aminosulfonyl)-7-fluoro-2, 1, 3-benzoxadiazole) in 200 mM sodium borate buffer pH 8.0 containing 1 mM EDTA and 0.1 volume equivalent (v/v) of 10 % tri-n-butyl phosphine in acetonitrile (13). The mixture was incubated in a 50°C water bath for 5 minutes and vortexed thoroughly after incubation. The mixture was placed on ice and 0.01 volume equivalent (v/v) of 2.5 N HCl added. The fluorescently labeled sample was used for injection on the chiral HPLC column and the ABD adducts detected using fluorescence. The areas under the respective peaks were quantified based on a standard curve of purified L and D cysteine.

### Western blot

Pancreatic and islet lysates were run on a 1 mm 4-12% Bis-Tris gel (Novex Life Technologies; Thermo Fisher; #NP0321) with MES SDS running buffer (Invitrogen; #NP0002-02) initially at 75 V and then at 120 V. The samples were then transferred to a prewet immobilon PVDF transfer membrane (Merck Millipore Ltd; #IPFL00010; Pore Size 0.45 μm) and sandwiched between wet filter paper and cassette holder. The entire apparatus was placed in a wet transfer apparatus (Bio-Rad) and run at 90 V for 90 min on ice. After transfer, the membrane was removed and incubated in blocking buffer containing Tris buffered saline + Tween 20 (TBS-T) containing 5% BSA for 30 minutes at room temperature (RT). The membrane was incubated on a shaker with the respective primary antibody at 1:1000 dilution in blocking buffer overnight at 4°C. After overnight incubation, the membrane was rinsed 3 times with TBS-T and washed in TBS-T at RT for 15 minutes per wash. The membrane was washed a total of 4 times. After washing, the membrane was incubated with HRP conjugated secondary antibody (IgG; mouse or rabbit) (GE Healthcare UK) at 1:5000 dilution in blocking buffer and incubated at RT with shaking for 1 h. After incubation with secondary antibody, the membrane was washed 4 times with TBS-T. The duration of each wash was 15 min. After the last wash, the membrane was incubated with Enhanced Chemiluminescent (ECL) reagent (Thermo Scientific; Super Signal West Pico Plus Peroxide solution (#1863099) and Luminol enhancer solution (#1863098) in a 1:1 (v/v) for 5 min in the dark. The excess ECL reagent was removed using a kimwipe and the membrane was placed in between plastic sheet in a cassette holder and developed on an 8” X 10” UltraCruz autoradiography film (Santa Cruz Biotechnology; #SC-201697) in a developer.

### Dot Blot of Secretory Vesicles and Exosomes

Dot blotting of secretory vesicles and plasma exosomes was performed on a prewet PVDF membrane treated with methanol, water and transfer buffer. The wet PVDF membrane was then placed in a 96 well microsample filtration manifold (Schneider & Schuell, NH) attached to a vacuum line. The samples were added to each well under suction. The suction was done till the samples were adsorbed on the membrane and the membrane dry. The microfiltration apparatus was then disconnected from the vacuum line and the PVDF membrane gently removed. The membrane with sample spots was then incubated in blocking buffer (5 % BSA+TBST) at room temperature with shaking for 30 minutes. Primary antibodies (rabbit insulin antibody: #4590 Cell Signaling Technology, OC (generic amyloid antibody) and rabbit β-actin antibody; #AC026 AB clonal) were added to the membrane at 1:1000 dilution and incubated overnight with shaking at 4°C. After incubation, the membrane was washed 4 times with TBST at RT for 15 minutes each. The membrane was then incubated with respective secondary antibodies at 1:5000 dilution in TBST at RT for 60 minutes. The blot was then washed 4 times with TBST at RT for 15 minutes each and developed using chemiluminescence with Enhanced Chemiluminescent (ECL) reagent (Thermo Scientific; Super Signal West Pico Plus Peroxide solution (#1863099) and Luminol enhancer solution (#1863098) in a 1:1 (v/v) for 5 min in the dark. The blot was exposed to X-ray film and developed.

### Isolation of Secretory Vesicles from βTC-6 cells

βTC-6 cells were grown in 30 cm dishes at 37°C, 5 % CO_2_, for 5-7 days in Dulbecco’s Modified Eagle’s essential medium (GIBCO, Grand Island, NY) supplemented with 15 % fetal bovine serum. Isolation of and purification of secretory granules from these cells were performed according to Hutton *et. al* (27) with slight modifications. Briefly, the cells were harvested and rinsed with PBS. After washing with PBS, the cells were resuspended with 1X hypotonic medium (2X medium: 60 % sucrose+ 100 mM EGTA-K+-1M MgSO_4_+ 100 mM MES in 50 ml water). The cells were homogenized in hypotonic medium using a Dounce homogenizer (Wheaton; Milville, NJ) with 20 strokes and on ice. The homogenate was centrifuged at 800 g for 5 min at 4C°. The supernatant was retained. The pellet was resuspended in original volume of hypotonic medium and centrifuged at 800 g for 5 min. The supernate was pooled from the first spin and centrifuged in a 13 x 56 mm polycarbonate centrifuge tube at 20,000 g at 4°C for 10 min in a Beckman SW55.TI rotor. The supernate was discarded and the pellet resuspended thoroughly in hypotonic medium. The pellet (500 μl volume) was then added to a new polycarbonate tube and underlayed with 3 ml isotonic 30 % Percoll mixed and centrifuged for 60 min at 25,000 g, 4°C in a SW55.TI rotor. After the spin, the gradient was fractionated and 200 μl aliquots sequentially discarded from the top till the last 800 μl was reached. The secretory vesicles were present in this fraction at the bottom of the tube. The secretory vesicle fraction was then centrifuged in the same tube for 10 min at 20,000 g and pellet stored at −80°C or used immediately for experiments.

### Isolation of plasma exosomes

Exosomes from mice plasma were harvested from age matched WT and SR^−/−^ mice (6-8 weeks) plasma using the total exosome precipitation reagent for plasma (Invitrogen #4484451) as per instructions. Plasma was isolated either on the day of exosome isolation or kept at −80°C and thawed for isolation of exosomes.

### Thioflavin T binding

Plasma exosomes and secretory vesicles from βTC-6 cells were assayed for β-sheet aggregates using ThT binding (33,34). All samples were assayed in quadruplicate with 10 μg of protein per well of a 96 well optical bottom microtiter plate (Thermo Fisher Scientific, Rochester NY). The total volume per well was 200 μl comprising 10 μg of exosome lysate or βTC-6 secretory vesicle lysate followed by ThT dye (final concentration of 1 mM) and PBS. The samples were mixed by pipetting and read immediately and subsequently at regular intervals at 11_ex_= 450 nm and 11_em_=482 nm. During the readings the plate was incubated at 37°C without shaking. The mean of the blank readings was subtracted from the mean of the sample readings at each time point and the corrected values, along with the mean and SD were plotted using KaleidaGraph (v4.1, Synergy Software, Reading, PA). Statistical analyses on the data (*t*-test) were performed using KaleidaGraph (33,34).

### Asc1 and ASCT2 Transport Studies

D-cysteine transport experiments were performed in HEK293 cells transiently transfected with human SLC1A15 (ASCT2; Origene; Cat# RC200305) and rat SLC7A10 (Asc1; Genomics-online.com; Cat# ABIN4047848) cDNA using lipofectamine 3000 reagent. Following 72 h incubation, the cells were harvested, washed twice with HEPES buffered saline (HBS) containing 10 mM HEPES and 100 mM NaCl, counted and equal number of cells (4 x10^7^ cells) distributed in tubes for each concentration of D-cysteine ranging from 0-120 μM. The cells were incubated with the different concentrations of D-cysteine at 37°C for 30 minutes. After incubation the cells were centrifuged at 3000 rpm for 5 min at RT and buffer discarded. The cells were washed 2 times with 500 μl HBS. The cells were hypotonically lysed by adding 200 μl cold sterile water to each tube and incubated at 37°C for 20 min. After hypotonic lysis 200 μl of reaction buffer (500 mM Tricine, 100 mM MgSO_4_ and 2 mM EDTA, 1% Triton X-100 pH 7.8) containing TCEP, CHBT and K_2_CO_3_ was added to each tube to perform the luciferase assay to measure D-Cysteine levels in the cells. The luciferase assay was performed as mentioned above and luminescence measured in the cell free supernates transferred to a 96 well opaque plate in a SpectraMax 96 well microplate reader and D-Cysteine levels quantified.

### Nuclear Extracts Preparation

Nuclear extracts were prepared from the pancreas of age matched 6-8 weeks old WT and SR^−/−^ mice as per instructions in the EpiQuik Nuclear Extraction Kit (Cat# OP-0002; Epigenetek, Farmingdale NY). The protein concentration of the samples was estimated using BCA protein assay and equal amounts of protein used in the DNMT activity assay.

### DNMT Activity Assay

Total DNMT activity was measured in nuclear extracts of WT and SR^−/−^ pancreas using the EpiQuik DNMT activity/inhibition assay colorimetric kit (Cat# P-3009; Epigenetek, Farmingdale NY). Equal amounts of nuclear extracts (20 μg) were added to each well of the 96 well plate and instructions followed as per the catalog. The activity of the nuclear extract was calculated as per instructions in the catalog and percent inhibition/activity plotted.

### Electron Microscopy of exosomes

Plasma exosomes were imaged using transmission electron microscopy. Samples (10 μl) were adsorbed to glow discharged (EMS GloQube) ultra-thin (UL) carbon coated 400 mesh copper grids (EMS CF400-Cu-UL), by floatation for 2 min. Grids were quickly blotted then rinsed in 3 drops (1 min each) of TBS. Grids were negatively stained in 2 consecutive drops of 1% uranyl acetate with tylose (UAT), and quickly aspirated to get a thin layer of stain covering the sample. Grids were imaged on a Hitachi 7600 TEM (or Philips CM120) operating at 80 kV with an AMT XR80 CCD (8 megapixel) camera.

### Methyl cytosine (mC) and Hydroxymethyl cytosine (hmC) DNA Dot Blot

DNA dot blot was performed on a nitrocellulose membrane. Briefly, 200 ng of DNA was mixed with 20X SSC buffer and Milli Q water. The sample was heated at 100°C for 10 min and cooled on ice. 200 μl of 20X SSC buffer was added to the sample. The nitrocellulose membrane was pre wet in SSC buffer and applied to a 96 well vacuum manifold and washed with 20X SSC buffer and samples applied to the 96 wells of the manifold under vacuum. After the samples were adsorbed on the membrane, the membrane was placed in denaturation solution (1.5 M NaCl + 0.5 M NaOH) for 10 minutes and in neutralization solution (1M NaCl + 0.5 M Tris-Cl pH 7.0) for 5 minutes. The membrane was partially dried and placed on a filter paper wet in 20X SSC buffer and UV crosslinked in a 2400 Stratalinker in auto crosslink mode. The membrane was then incubated in blocking buffer for 30 minutes at RT with gentle shaking. After blocking the membrane was incubated with rabbit mC and hmC antibodies (1:2000 dilution) in TBST overnight at 4°C. The membrane was washed four times 15 min each in TBST and incubated with secondary antibody (1:5000 dilution) in TBST for 1 h at RT. The membrane was developed using ECL chemiluminescence on an X ray film.

### cAMP Live Cell Imaging

Imaging of live HEK293 and βTC-6 cells was performed on a 3i spinning disk confocal microscope with slide chamber set up maintained at 37°C and 5 % CO_2_ to maintain pH and cell viability. Image settings and parameters were controlled using Slidebook 6 software. Cells were seeded prior to imaging and transfection in 35 mm dishes with in built coverslip. Imaging was performed at 20X magnification and over 5 minutes duration with 30 second time lapse. Filter set was c562mp. For each field, 8-10 points were highlighted in the XY plane for capture. These points were based on maximal intensity of cell fluorescence in the active viewing field. Such points were highlighted in multiple fields of view. For time-lapse capture, the 8-10 points were captured at 30 second intervals for a total duration of 5 minutes. The fluorescence intensities were obtained from Image J, background subtracted and plotted versus elapsed time. Movie files were generated in Image J by combining each individual live capture file [42].

### Chromatin Immunoprecipitation-qPCR (ChIP-qPCR)

ChIP was performed from pancreas of age matched WT and SR^−/−^ mice using EpiQuik tissue chromatin immunoprecipitation kit (Epigentek; #P-2003) as per instructions. Rabbit anti CREB monoclonal antibody (Cell Signaling Technology; #D76D11) was used as per recommended dilution (1:100) to IP DNA. Following DNA elution and quantitation, 100 ng of DNA was used in qPCR reactions using the SYBR green method. Input DNA C_T_ values was used to subtract the C_T_ values from WT and SR^−/−^ samples and fold change relative to WT was determined using 2^-ΔΔCT^. IgG antibody was used as a negative control. DNMT1 promoter region (D1-D5) primer sequences used for qPCR are listed below [47].

D1 Reverse: 5’ AAC GAG ACC CCG GCT TTT T 3’

D1 Forward: 5’ TAT AGC CAG GAG GTG TGG GTG 3’

D2 Forward: 5’ TCC TCT GCA AGA GCA GCA CTA 3’

D2 Reverse: 5’ ATG TAC CAC ACA GGG CAA GA 3’

D3 Forward: 5’ TGT TTG TGC ATG TGA GTG CA 3’

D3 Reverse: 5’ TCG GCA CTT GAG AGC AGG TA 3’

D4 Forward: 5’ TGA GTG CTG GAA TCA AAT GC 3’

D4 Reverse: 3’ AAG CCC CTG TAA TTC CAC TT 3’

D5 Forward: 3’ AGA AGT GGT TCC TGG CCT TA 3’

D5 Reverse: 3’ TAA CTC TAT CCC CCT CCC CTT 3’

### Bioinformatic Analysis

Proteins identified in the proteomics screen were annotated and classified into pathway and as gene ontology (GO) as overexpressed (n=196) and under expressed (n=33) in SR^−/−^ when compared to WT. Protein enrichment, annotations, functional pathway and GO analysis were performed using ShinyGO 0.76 (http://bioinformatics.sdstate.edu/go/) which converts the Protein IDs to their corresponding Gene IDs.

### Glucose Stimulated Insulin Secretion (GSIS) in mouse islets and INS-1 832/13 cells

Mouse islets and INS-1 832/13 cells secrete insulin in media in response to glucose and other secretagogues. GSIS was performed in these cells (passage >4) in response to 10 mM (final concentration) D-cysteine and D-serine at 37°C along with 100 mM glucose. Briefly, 0.5-1×10^6^ cells were plated in each well of a 12 or 24 well plate in basal glucose media (66) and allowed to reach 75 % confluence (approx. 24-48h). The media was changed to glucose free media and D cysteine and serine were added to individual wells. For both time course and dose response, the cells were treated with D-serine and or D-cysteine alone in absence of glucose. For both these experiments, glucose alone treatment (100 mM) served as a positive control. For islet experiments, the isolation was performed as per the ref and the concentrations of the secretagogues were kept at 10 mM [68]. The media was harvested in a tube containing 1X protease inhibitor cocktail and stored in −80°C. The media was subsequently assayed for insulin secretion using rat insulin ELISA kit (proteintech; cat no: KE20008 96T), mouse insulin ELISA kit (for mouse islets; proteintech; cat no: KE10089) and human insulin ELISA kit (crystal chem; #90095). Islet viability was assessed by propidium iodide and fluorescein di acetate staining. Insulin concentration was estimated using the respective standard curve. Data are representative of 2 independent experiments. The data and statistics (student’s *t-test*) were plotted using Kaleidagraph (Synergy Software). Western blot on INS-1 832/13 cells was performed at different doses of D-serine and D-cysteine with T=6 h incubation at 37°C. Glucose at 200 mM was a positive control.

### Streptozotocin (STZ) injection in C57BL/6J mice

8-12 weeks old male C57BL/6J mice were intraperitoneally injected with STZ at 40 mg/kg body weight once daily for 10 days. On days 2, 4, 6, 8 and 10 of STZ administration, mice were euthanized by CO_2_ narcosis followed by cervical dislocation and plasma and pancreas harvested.

### Human Islet experimentation

Normal human islets were purchased from Prodo Labs Inc (Aliso Viejo, CA). A total of 1000 islet equivalents was purchased. Purity of the islets was 95 % and viability was 95 %. The islets were centrifuged at 350 g for 5 minutes and resuspended in 4 ml of CMRL medium. The resuspended islets were equally distributed in a 12 and 6 well plates with each well containing 50 islets. For GSIS, the concentration of glucose was 10 mM. The concentrations of D-serine and D-cysteine was 10 mM.

## 6. Acknowledgements

The authors acknowledge Ms. Barbara Smith of the Microscopy Core Facility at Johns Hopkins School of Medicine for assisting in laser scanning confocal microscopy, live cell imaging and obtaining electron micrographs of secretory vesicles and exosomes. The authors acknowledge Evan R. Semenza for helpful discussions and Ms. Susan McTeer for her administrative assistance. The authors also acknowledge MMG and MA who tragically passed away in 2018 and 2022 respectively.

## 7. Author contributions: CRediT

Conceptualization: RR

Methodology: RR,

Reagents: SHS, MA, PY

Investigation: RR, TW, HS, LA, SB

Visualization: RR

Funding acquisition: SHS

Project administration: RR

Supervision: RR

Writing: RR

Writing – review & editing: RR, SHS.

## 10. Data and materials availability

All data are available in the main text or the supplementary materials and available on request.

