## Supplementary figures and images for "Mammalian D-Cysteine controls insulin secretion in the pancreas"

### Figure S1

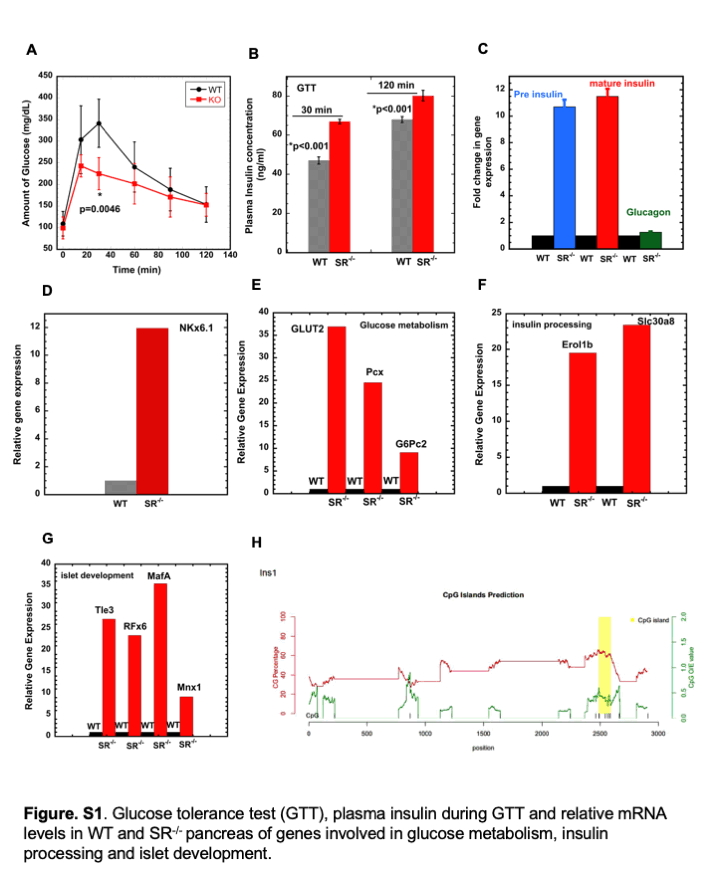

### Figure S2

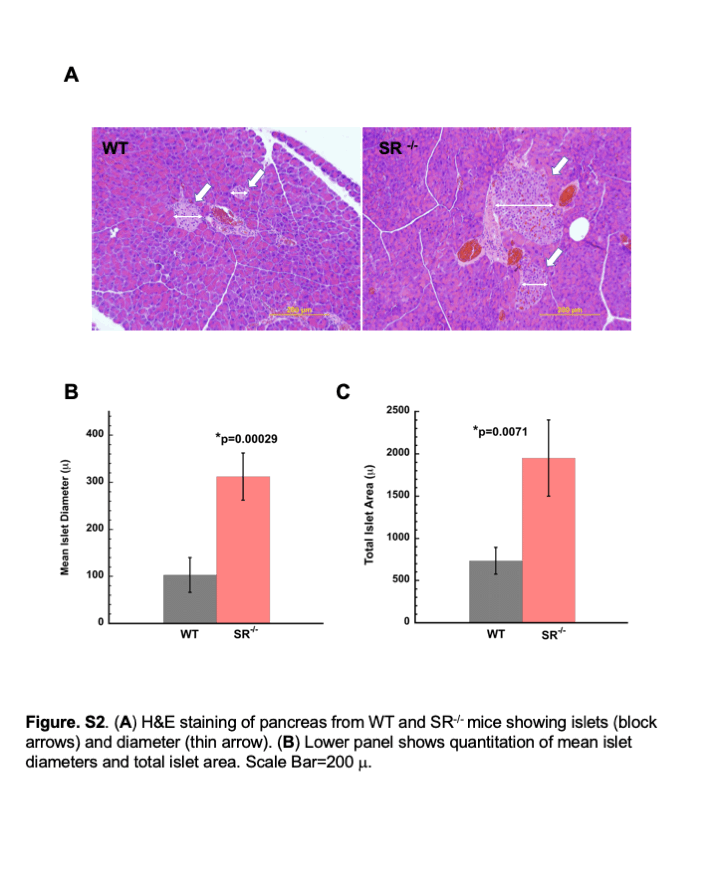

### Figure S3

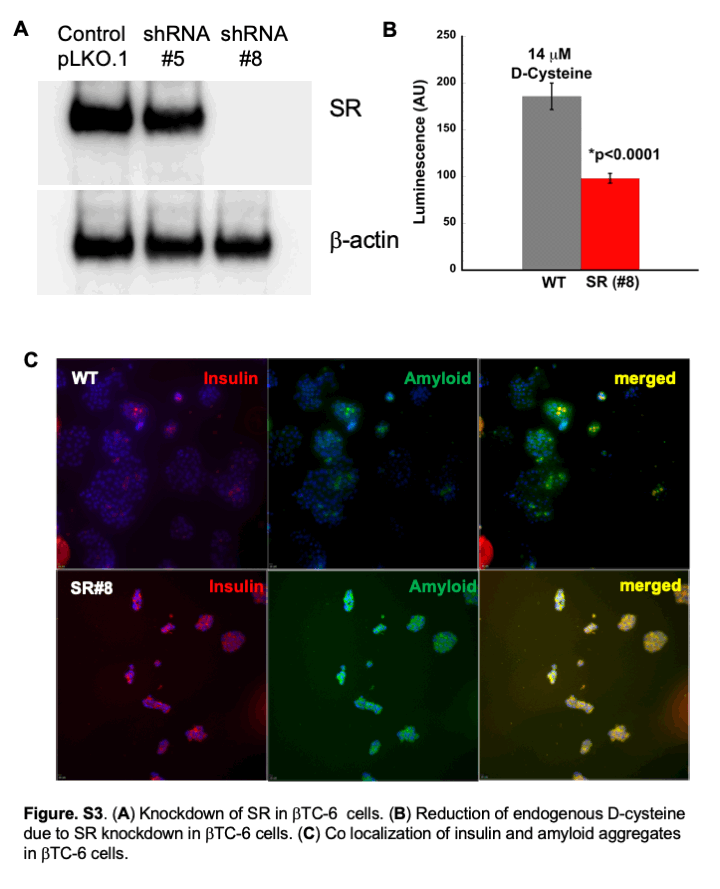

### Figure S4

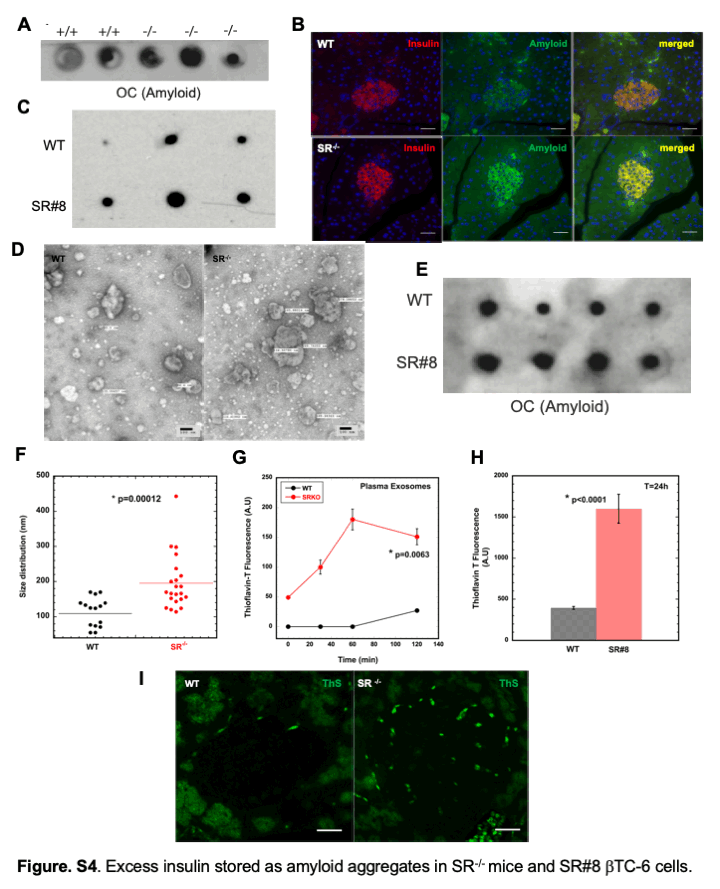

### Figure S5

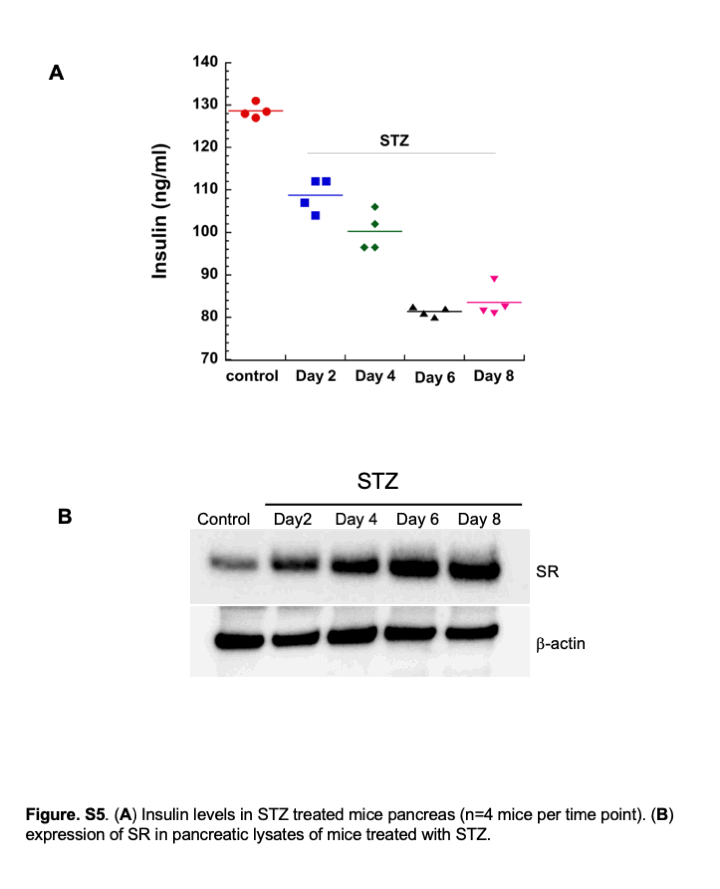

### Figure S6

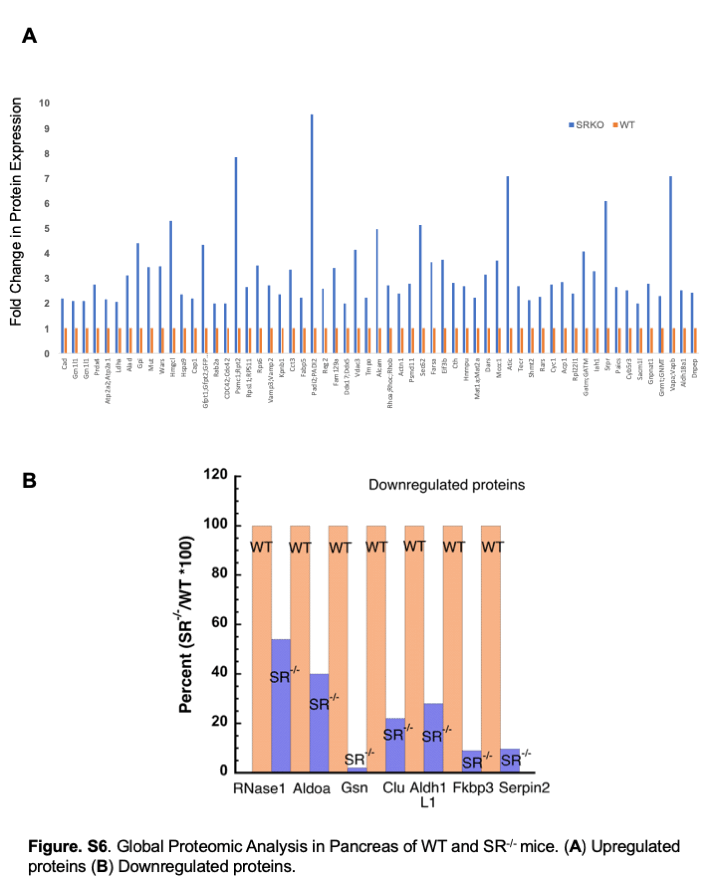

### Figure S7

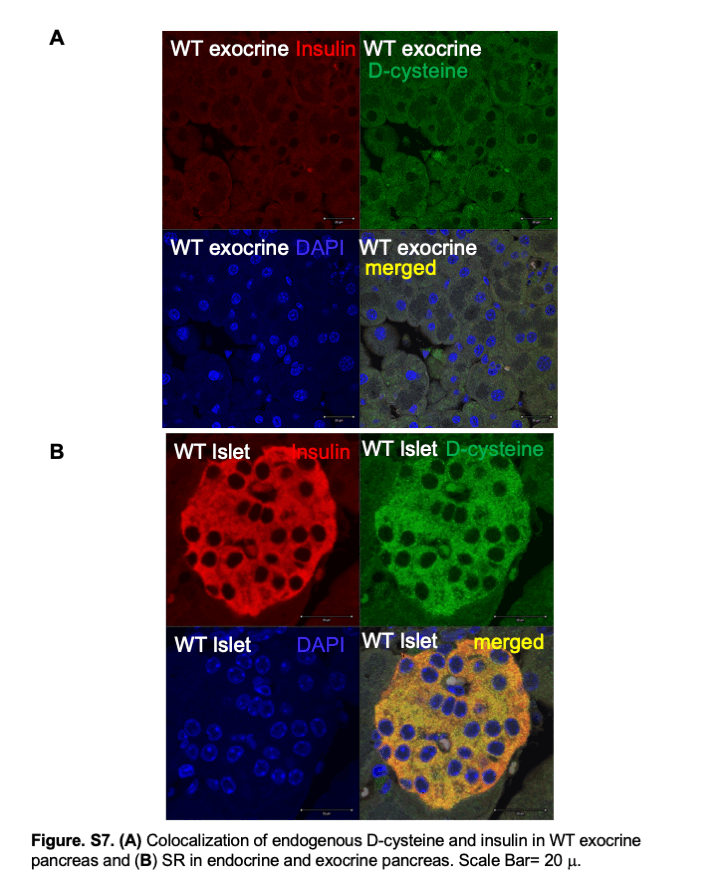

### Figure S8

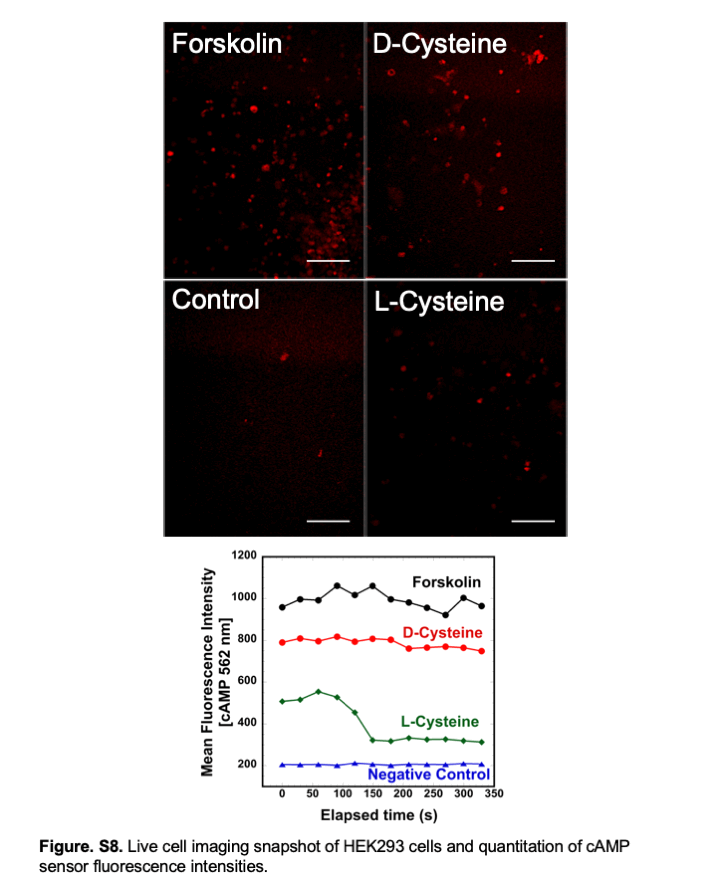

### Figure S9

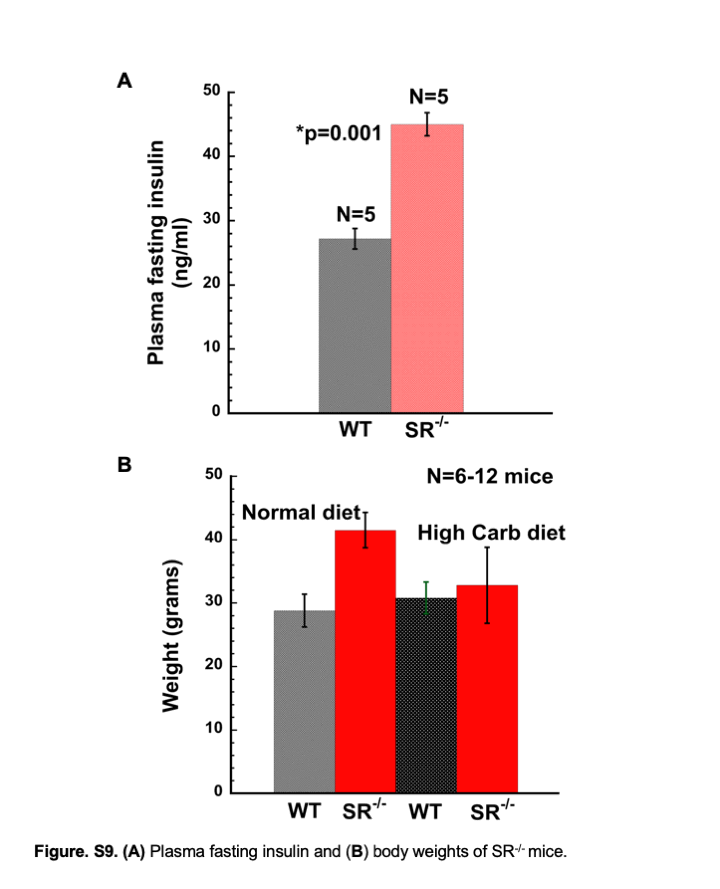

### Figure S10

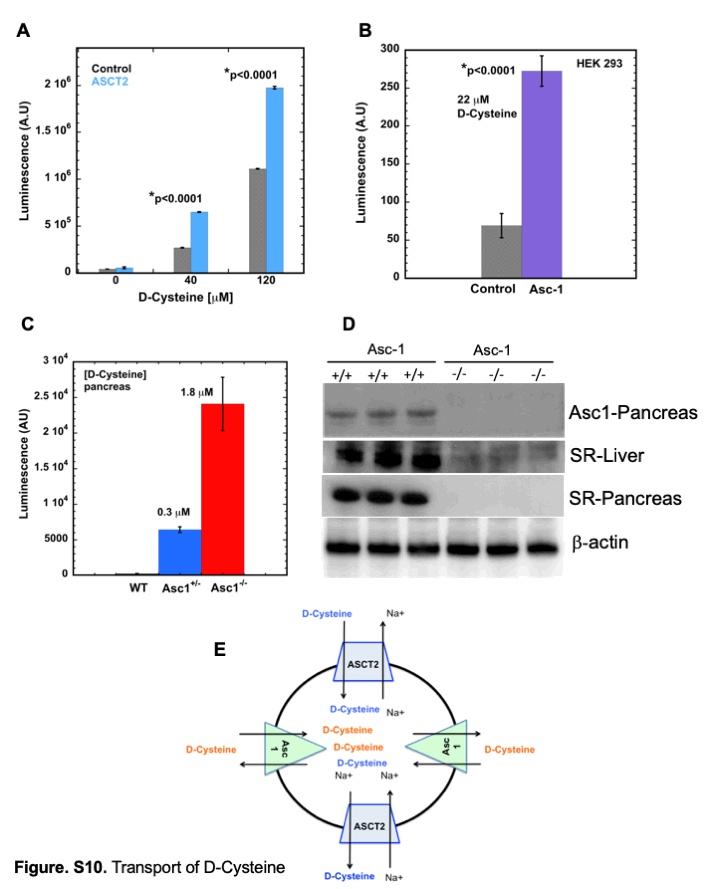

### Table 1

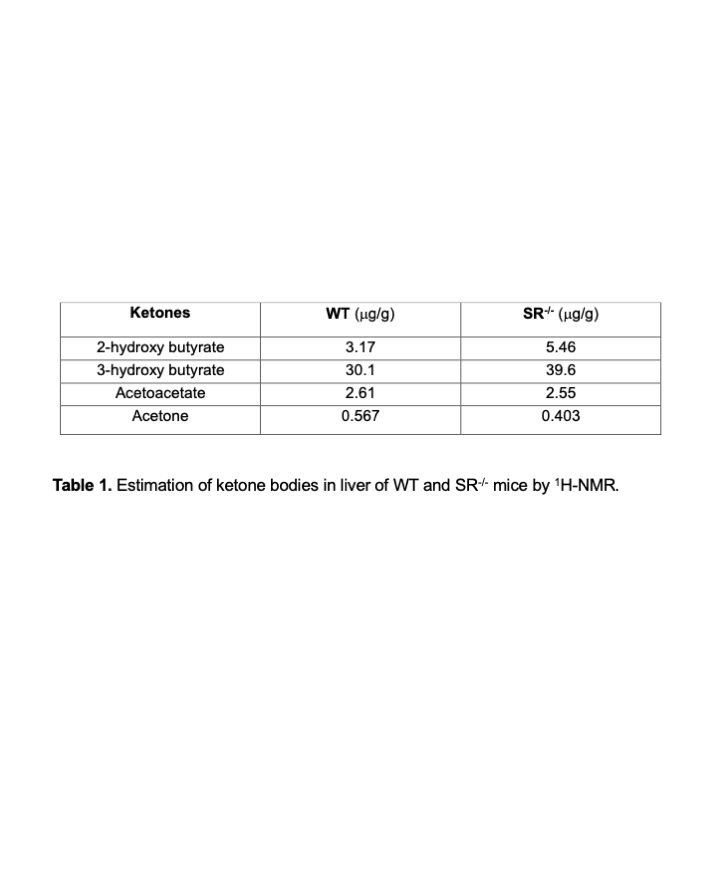

### Table 2

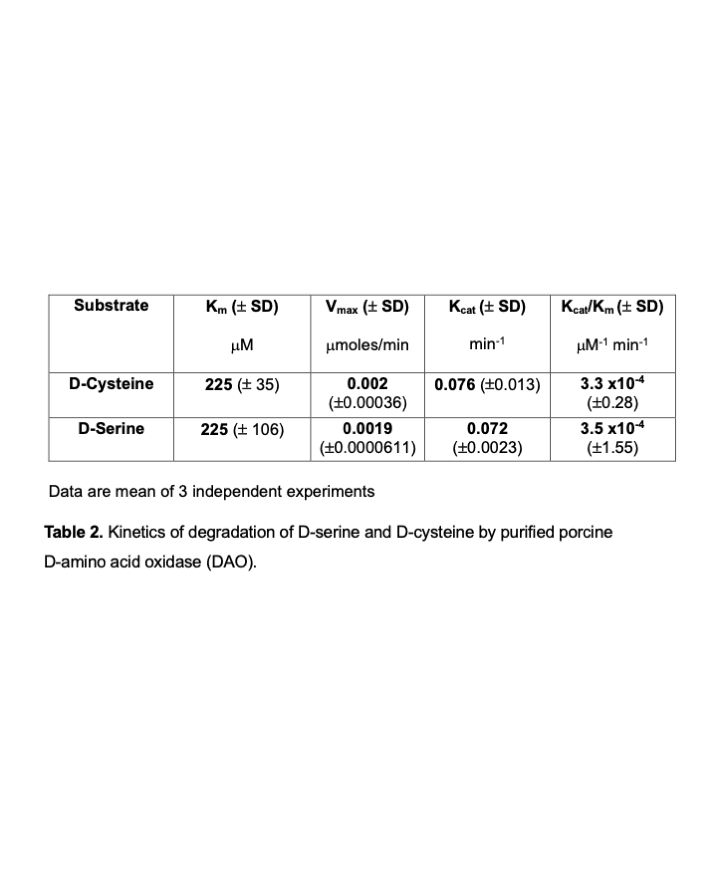

### Table 3

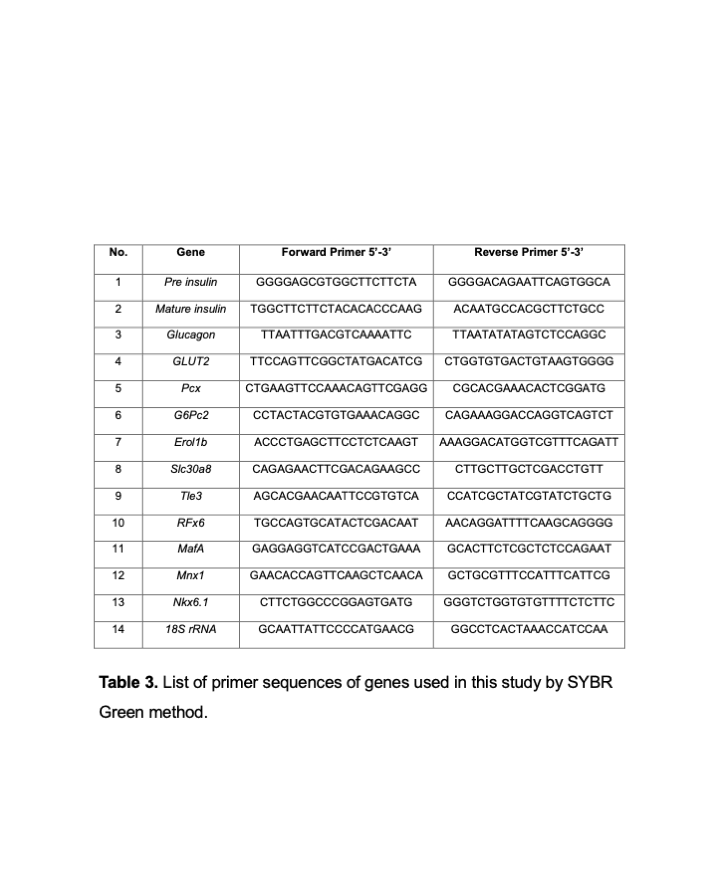

### Table 4

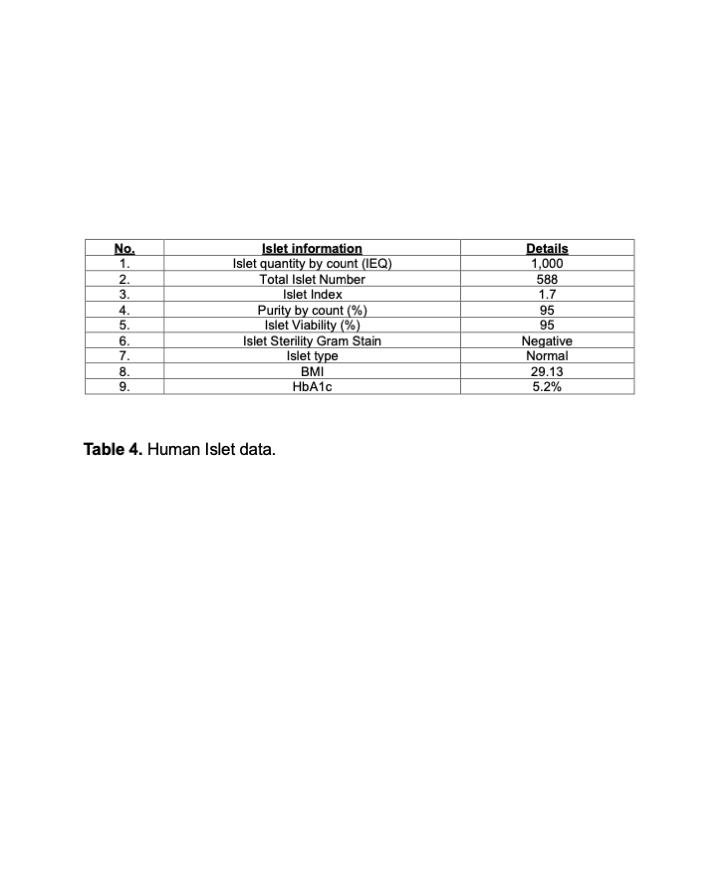
